# Nutrient transporter translocation to the plasma membrane via a Golgi-independent unconventional route

**DOI:** 10.1101/540203

**Authors:** Vangelis Bouris, Olga Martzoukou, Sotiris Amillis, George Diallinas

**Author notes:** Equal contribution (Vangelis Bouris, Olga Martzoukou).

## Abstract

Nutrient transporters are believed to traffic from their site of synthesis, the ER, to the plasma membrane, through the Golgi, using the conventional vesicular trafficking pathway. However, here we report that the UapA purine transporter in *Aspergillus nidulans*, follows a Golgi-independent, unconventional new route, which does not involve key Rab GTPases, AP adaptors, microtubules or endosomes, but is dependent on functional COPII, clathrin heavy chain (ClaH) and actin. The role of ClaH in transporter secretion is shown to be unrelated to that performing, together with Rab11/AP-1, at the Golgi, and is seemingly due to an effect in actin network functioning. Our findings are discussed within the frame of a model that rationalizes why the trafficking mechanism uncovered herein might hold true for transporters in higher organisms.

**One Sentence Summary:** Fungal transporters use a non-polar, COPII- and actin-dependent, route for subcellular traffic to the cell membrane.

## Main Text

Plasma membrane (PM) transporters mediating the selective cellular uptake or efflux of solutes and drugs are essential proteins in all organisms. The first step in their biogenesis, being polytopic transmembrane proteins, is their co-translational translocation into the membrane of the Endoplasmic Reticulum (ER). The current belief is that after translocation into the ER, transporters are sorted into nascent ER-exit sites, pack into budding COPII vesicles that fuse to the cis-Golgi and then reach the *trans-Golgi* network (TGN) via Golgi maturation *(1–3).* From the TGN, transporters are thought to be secreted towards the PM, similar to other membrane cargoes, either directly or indirectly via the endosomal compartment, in AP-1/clathrin coated vesicles, the trafficking of which is controlled by multiple Rab GTPases and the microtubule cytoskeleton *(4, 5).*

However, some lines of evidence support that *de novo* made transporters, such as the insulin-regulated GLUT4 glucose carrier, might not follow known conventional Golgi and post-Golgi dependent routes. For example, genetic knock-out of proteins involved in TGN-dependent membrane cargo sorting (e.g. Arfrp1, golgin-160 or AP-1) leads to GLUT4 accumulation in the PM, rather than retention in the Golgi or other intracellular compartments, suggesting the presence of an unknown Golgi-independent route *(6).* In addition, kinesin motor proteins or microtubule disruption have a moderate or no effect on GLUT4 accumulation at the PM *(7, 8).* In line with evidence concerning GLUT4, no formal evidence exists on whether several other *de novo* synthesized transporters traffic through the Golgi/TGN compartment. A possible explanation for this might be that PM transporter passage and exit from the Golgi is very rapid, never leading to accumulation of sufficient steady state levels for detection with standard fluorescence microscopy. However, evidence against this explanation is also the fact that no mutation or specific condition has been shown to block PM transporters exclusively in the Golgi. Instead, several mutations affecting the proper folding or specific motifs in transporters are known to lead to retention in the ER, often associated with ubiquitination-dependent turnover by proteasome degradation and/or selective autophagy *(9, 10).* Importantly, the CTFR transmembrane protein, an ATP-binding cassette (ABC) transporter that functions as a low conductance Cl^-^ selective channel associated with cystic fibrosis, has been formally shown to translocate to the PM via direct trafficking from the ER in COPII vesicles *(11).*

In this work we provide evidence that nutrient transporters in the fungus *Aspergillus nidulans* are sorted in the PM following a Golgi-independent trafficking route, involving COPII-dependent exit from the ER, a functional clathrin heavy chain, and proper acting network organization. We believe that the exocytic route identified herein might well be the principle mechanism by which nutrient transporters, and probably other non-polar house-keeping membrane cargoes are targeted to the PM in also in animals or plant cells.

## Results

### Sorting of newly made UapA from the ER to the PM

To investigate whether newly made PM transporters follow the conventional secretory pathway involving the Golgi and other known post-Golgi trafficking effectors, we constructed two strains allowing controllable expression of *de novo* made GFP-tagged functional version of the well-studied uric acid-xanthine transporter UapA*(14, 15).* One strain expresses UapA via its native promoter, whereas the second uses the rather stronger *alcAp* promoter *(15).* These strains, named UapA or *alcAp-* UapA, both allow transcriptional repression-derepression of UapA at any moment during *A. nidulans* conidiospore germination and vegetative growth of hyphae (for details see Materials and methods). In order to follow sorting to the PM of neosynthesized UapA, we allowed *A. nidulans* conidiospores to germinate overnight *(12–14)* under conditions which transcriptionally repress UapA synthesis, and then induced its *de novo* synthesis. We thus followed the subcellular localization of newly made UapA in young hyphal cells, by *in vivo* fluorescent microscopy at 0-4 hours after derepression of transcription. Results in **Fig. 1A** and **1B** show that no or very little UapA-GFP fluorescence is detected under repressed conditions(0 h),while upon imposing de-repressing conditions UapA appears first in a cytosolic membranous mesh (80-100 min), later in more cortical membranous and punctuate structures (100-120 min), and finally in the PM (140-180 min). When UapA is expressed via the *alcAp* promoter it also labels characteristic ER perinuclear rings (see the 100-120 min panel in **Fig. 1B**), and is generally expressed at higher levels (notice that PM labeling at 180-190 min appears homogeneous, compared to the rather punctuate labeling when expressed from the native *uapA* promoter). In general, however, the timing of secretion and subcellular localization of *de novo* expressed UapA is similar, irrespectively of the promoter used.

**Fig. 1.**
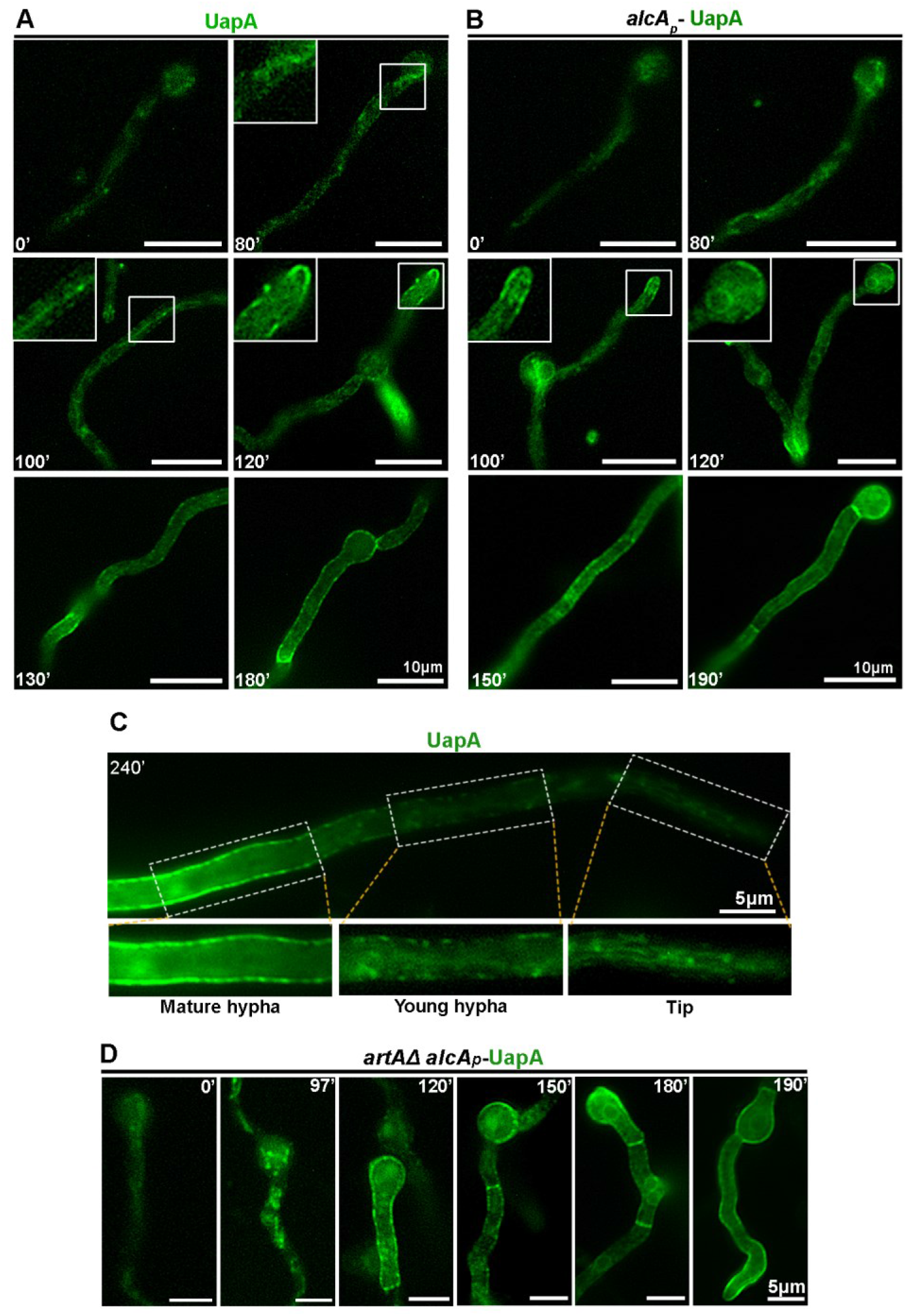
Subcellular localization of neosynthesized UapA. **(A)** Epifluorescence microscopy following *de novo* expressed UapA-GFP, in the presence of a non-repressive nitrogen source (NO_3_^-^), for 0, 80, 100, 120, 130 and 180 minutes. Notice that localization of a fraction of UapA at the PM is first observed after 120 min of derepression. **(B)** Similar experiment as in (A) following newly made UapA-GFP expressed via the *alcAp* promoter in the presence of non-repressive carbon (fructose) and nitrogen (NO_3_^-^) sources, for 0, 80, 100, 120, 150 and 190 min. Notice the apparent localization of the transporter in the PM and in perinuclear ER rings after 100-120 min of derepression. **(C)** Epifluorescence microscopy of a maturing hypha after 240 min of derepressed UapA-GFP expression. Notice that in the growing hyphal tip UapA localizes in a membranous mesh and some cytosolic puncta, whereas in more tip-distal parts of the hypha UapA is in PM-localized puncta, which become progressively abundant so that at more posterior areas the PM is homogeneously labeled. **(D)** Time course following UapA sorting to the PM under inducing conditions imposed for 0, 97, 120, 150, 180 and 190 min, in a genetic background where ArtA, the arrestin required for UapA endocytosis, is genetically depleted.

We also followed the spatial localization of *de novo* made UapA in longer, more mature, hyphae (240 min). We observed that while in older compartments UapA was exclusively and homogenously translocated in the PM, in younger parts, and especially close to the apical region, UapA was still detectable in a cytosolic membrane network and several, little-motile, foci (**Fig. 1C** and video in **Movie S1**). This shows that UapA is not translocated to the apical-distal PM via later diffusion from the tip area, as some apical cargoes do *(16).*

To confirm that what we observed corresponds to *bona fidae* secretory/exocytic localization of *de novo* made UapA, and exclude that part of the picture obtained is also due to UapA endocytosis and recycling from the PM, we repeated the same experiment in a strain expressing UapA-GFP in a genetic background defective for UapA endocytosis, due to a deletion of the arrestin adaptor ArtA that is necessary for HECT-type ubiquitination preceding UapA internalization from the PM *(17).* **Figure 1D** shows that we obtained a similar result, showing again the transient localization of UapA in a membranous network and the absence of Golgi-like bodies, before the transporter reaches its final PM destination. The noninvolvement of endocytosis and recycling in the picture obtained was further supported by the observation that transient UapA cytosolic structures are not colabeled with FM4-64, a molecular marker specific for endosomal membranes (**Fig. S1**). Thus, the overall picture is that *de novo* made UapA, before reaching the PM, localizes in a membranous network, apparently the ER, and in several little-motile cytosolic bodies, but in no case labels structures resembling Golgi bodies or endosomes.

### UapA trafficking from the ER to the PM is Golgi-independent

We then examined whether *de novo* made UapA translocates to the PM when a series of key Golgi regulators necessary for conventional secretion are transcriptionally knocked-down through the use of a thiamine-repressible *thiA* promoter (*thiAp*) *(15, 18).* The Golgi proteins tested for their role in UapA trafficking were: SedV^Sed5^, HypB^Sec7^, GeaA^Gea1^, RabC^Rab6^ and RabO^Rab1^ *(19).* In brief, SedV is an early Golgi syntaxin, HypB and GeaA are Golgi guanine nucleotide exchange factors, and RabC and RabO are small regulatory GTPases. These proteins are all present across the Golgi and though it is difficult to define their exact site of action, SedV and GeaA are considered early (ER-Golgi interface) to medial Golgi regulators, RabO and RabC are medial-late Golgi regulators, while HypB is a late-TGN regulator. All these proteins are essential for Golgi functioning and apical cargo trafficking *(20, 21).* This is in line with the observation, that when we repressed transcription of the corresponding genes, vegetative growth of *A. nidulans* was arrested or very much reduced, after initial germination (**Fig. 2A**). Efficient thiamine-repression of the relevant Golgi genes is directly confirmed by western blot analysis, which depicts a > 90-95% reduction of steady state levels of the relevant proteins (**Fig. 2B**).

**Fig. 2.**
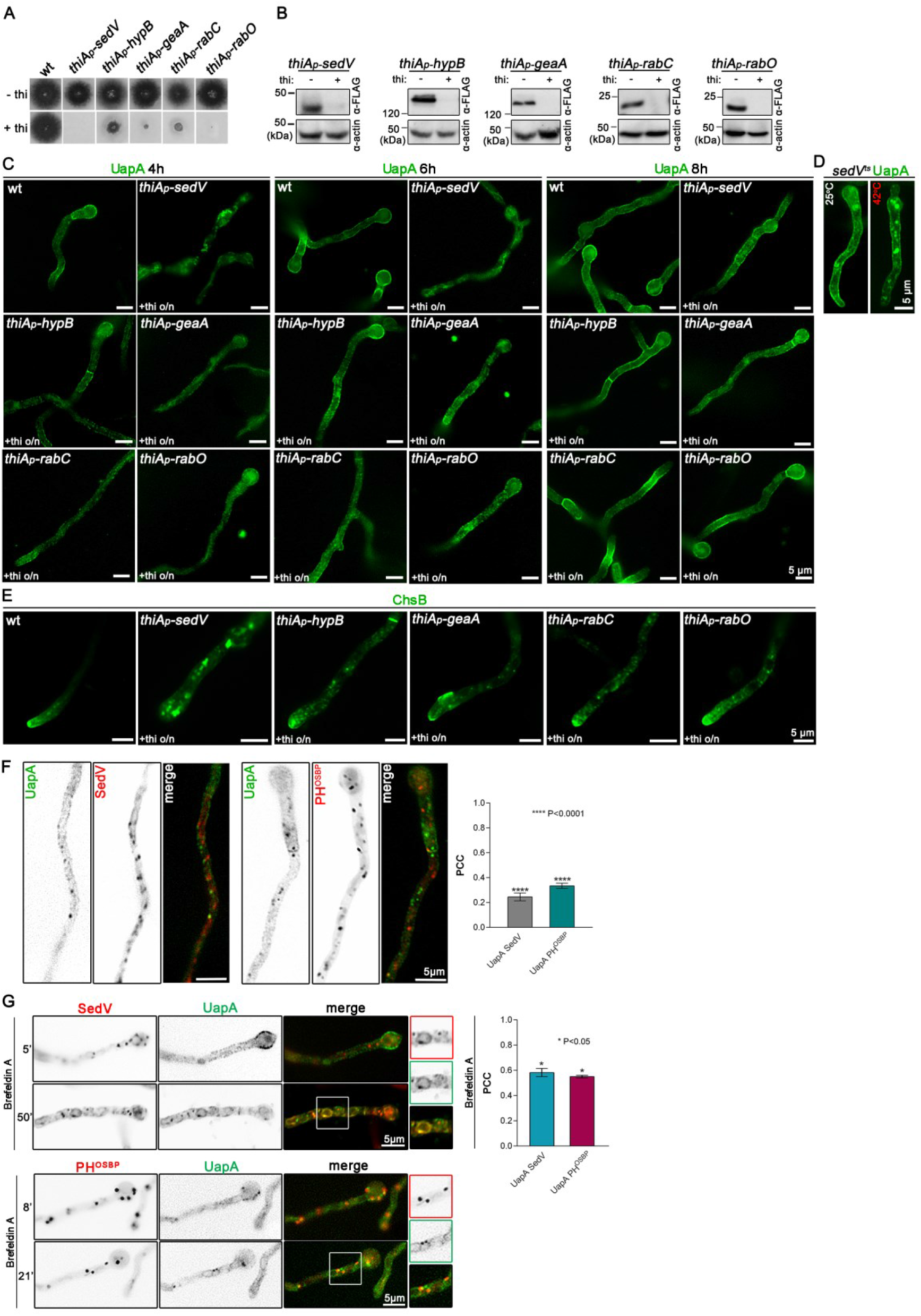
UapA translocation in the PM is Golgi-independent. (A) Growth of isogenic strains carrying thiAp-repressible alleles of *sedV, hypB, geaA, rabCand rabO* (*thiAp-sedV*, *thiAp-hypB, thiAp-geaA, thiAp-rabC*and *thiAp-rabO*) compared to wild-type (wt) in the absence (-) or presence (+) of thiamine. (B) Western blot analysis comparing protein steady state levels of FLAG-tagged versions of SedV, HypB, GeaA, RabC and RabO, in the absence (-) or presence (+) of thiamine, added *ab initio* and for 18 h (overnight culture). Equal protein loading is indicated by actin levels. (C) Microscopy analysis of *de novo* made UapA-GFP localization, expressed via derepression of the *alcAp* promoter, for 4, 6 or 8 h, under conditions where SedV, HypB, GeaA, RabC and RabO are *ab initio* repressed by thiamine. Notice that UapA reaches the PM in all cases, but the rate of translocation in the PM is delayed, in several cases, compared to the isogenic wild-type (Sec24 is required for proper ER-exit of UapA.wt) control. (D) Subcellular localization of *de novo* made UapA *(alcAp,* 150 min derepression) in a SedV^ts^ thermosensitive mutant (25 or 42 ^o^C for 90 min, after 60 min of derepression). Notice that even at the restrictive temperature (42 ^o^C) UapA reaches the PM. (E) Subcellular localization of the *de novo* made chitin synthase GFP-ChsB, in strains where SedV, HypB, GeaA, RabC and RabO are *ab initio* repressed by thiamine, compared to the wild-type (wt). Notice that in all cases proper localization of ChsB at the fungal tip is impaired (less so in the absence of RabC and RabO). (F) Co-localization analysis of neosynthesized UapA-GFP, prior to PM translocation (90 min derepression, *alcAp* promoter), with early (SedV-mCherry) and late (PH^OSBP^-mRFP) Golgi markers (left and middle panels, respectively). Notice the practically absent overlapping fluorescent signal with SedV (PCC=0.25, P<0.0001, n=4) or with PH^OSBP^ (PCC=0.34, P<0.0001, n=5). **(G)** Treatment of strains co-expressing SedV-mCherry or PH^OSBP^-mRFP and UapA-GFP (from *alcAp,* 90 min derepression) with the Golgi inhibitor Brefeldin A (BFA), for the time indicated. Notice that BFA elicits accumulation of early and late Golgi puncta in larger aggregates or perinuclear rings (in the case of SedV-mCherry). Quantification of UapA co-localization with SedV and PH^OSBP^ was not significant (PCC=0.58, P=0.0032, n=3 and PCC=0.55, P=0.0116, n=2, respectively) and seems to be the result of collapse and fusion of cis-Golgi membranes with ER.

When subcellular localization of *de novo* made UapA was followed for a period of 4-8 h following derepression of *uapA* transcription, under pre-established repression of the selected Golgi genes, we noticed that in all cases UapA translocates to the PM, similarly to the picture obtained in a wild-type background. However, in most cases, and especially when SedV was repressed, UapA secretion was delayed, as it took 8 h instead of 4 h of *uapA* derepression for the transporter to label massively the PM (**Fig. 2C**). Repression of the other Golgi proteins led to a more moderate delay, as already at 6h most of UapA labeled the periphery of cells. The delay in UapA sorting to the PM was very probably a consequence of the dramatic effect caused by the absence of Golgi functioning that is also reflected in the absence of development of micro-colonies, as shown in **Fig. 2A**. In the case of SedV, we additionally made use of a previously studied thermosensitive mutant SedVR^258G^ that allows a more immediate inactivation of SedV, and thus Golgi functioning, in the non-permissive temperature *(22).* As shown in **Fig. 2D**, shift of the SedVR^258G^ mutant at 42°Cdid not block effective translocation of *de novo* made UapA to the PM, despite a degree of degradation, reflected as cytosolic structures corresponding to vacuoles, caused by exposure to high temperature.

To confirm that the conditions used to follow UapA secretion lead to efficient knockdown of Golgi functioning, we also followed the secretion of ChsB chitin synthase. **Fig. 2E** shows that *ab initio* repression of *sedV, hypB geaA,* and to lesser degree of *rabC* and especially *rabO,* led to significant mislocalization of ChsB from its normal apical localization observed in a wild-type background, in agreement with previous reports *(20, 21).*

To obtain further evidence that *de novo* made UapA localizes to the PM via a Golgi-independent mechanism, we also performed co-localization studies with early (SedV) or late/TGN (PH^osbp^) Golgi molecular markers *(23).* We found, statistically supported (Pearson correlation coefficient (PCC) < 0.35), non-significant colocalization of UapA with either SedV or PH^osbp^ (**Fig. 2F**). We also examined whether UapA is trapped into Golgi aggregates, marked with SedV or PH^osbp^, obtained in the presence of Brefeldin A (BFA), a drug that leads to Golgi collapse and arrest in polar cargo trafficking *(23).* Although this approach led to a somehow increased correlation (PCC 0.55-0.58) of common localization of UapA and Golgi aggregates (**Fig. 2G**) this is judged non-significant, and seems to be due mainly to a negative effect of prolonged exposure to BFA on ER. An effect of BFA on ER organization has been reported before and shown to be due to the collapse of *cis*-Golgi membranes and fusion with those of the ER. This can be clearly seen after 50 min exposure to BFA SedV marks perinuclear rings typical of the ER.

### UapA exits the ER from ERes

We asked whether exit from the ER occurs via conventional ER cargo exit sites (ERes) by testing whether UapA secretion is blocked upon controllable knockdown of Sec24, a component of COPII-coated vesicles that mediates cargo vesicular exit from the ER. **Figure 3A** shows that, as might have been expected, thiamine-repression of *sec24* leads to a lack of growth of *A. nidulans.* **Figure 3B** shows that UapA secretion to the PM absolutely requires the expression of Sec24, as UapA remains massively in cytosolic structures that resemble ER aggregates, when transcription of the *sec24* gene is *ab initio* repressed. This block is persistent, as UapA never translocates to the PM, even after prolonged expression (12 h). A similar Sec24-dependent total block was observed when we followed, as a control, the secretion of an apical cargo, namely chitin synthase ChsB (**Fig. 3C**). We also tested whether UapA and Sec24 interact, using a co-localization approach. Despite the partial functioning of Sec24-mRFP (see **Fig. 3A**), the appearance of Sec24-specific puncta was typical of ERes *(23)* and colocalized significantly with UapA-labeled ER membranes (**Fig. 3D** – video in Movie S2). Unexpectedly, the co-localization experiment also showed that UapA secretion is somehow delayed in the strain expressing mRFP-tagged Sec24 (**Fig. 3D**). This delay is reflected in the prominent appearance of UapA-labeled perinuclear ER rings, not seen in an untagged Sec24 background. Thus, the partial functioning Sec24-mRFP fortuitously further confirmed that Sec24 is essential for proper UapA secretion.

**Fig. 3.**
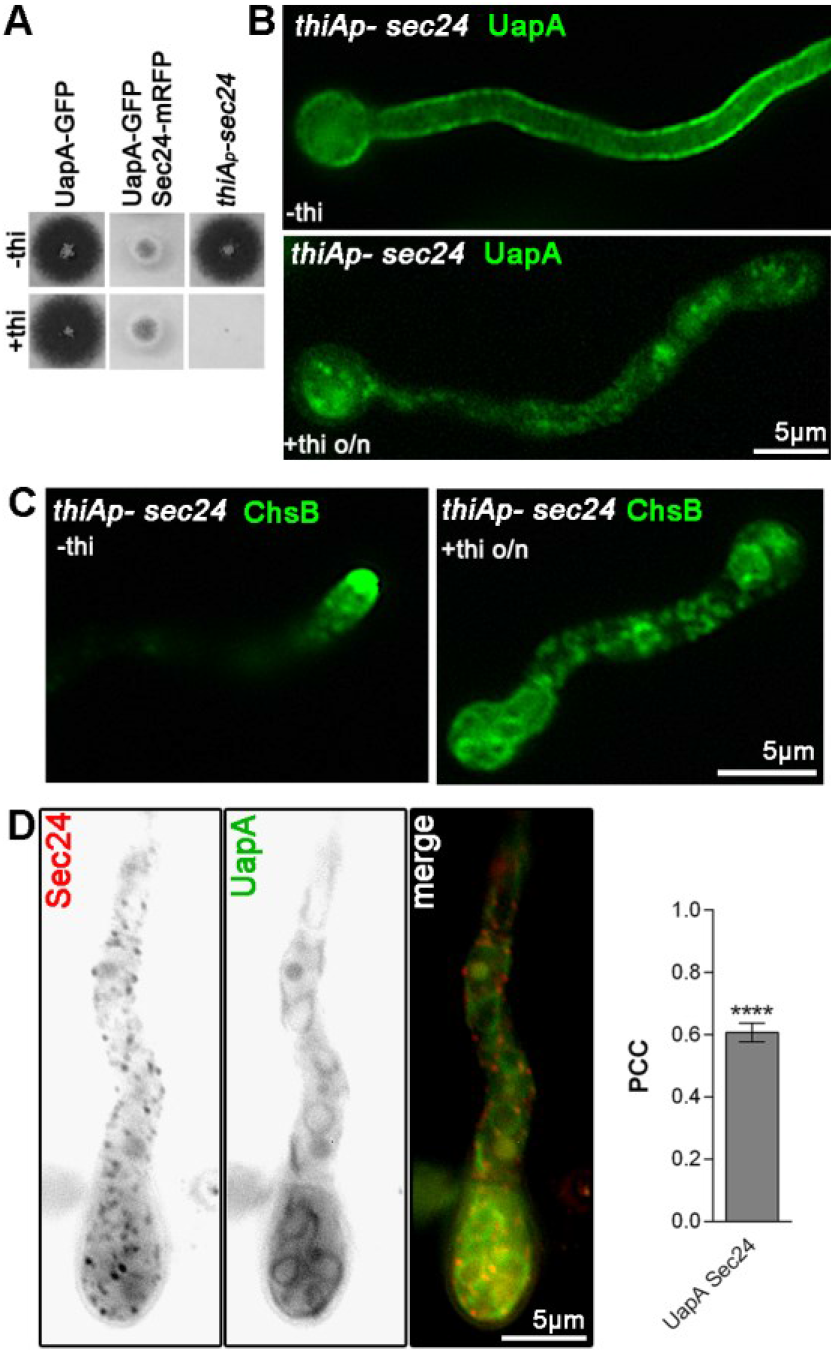
Sec24 is required for proper ER-exit of UapA. (A)Growth tests of isogenic strains showing the tagged versions of UapA and UapA/Sec24, compared to a strain carrying the *thiAp-sec24* allele, in the absence (-) and presence (+) of thiamine. (B, C) Epifluorescence microscopy analysis of the subcellular localization of UapA-GFP (B) or GFP-ChsB (C), under conditions where Sec24 expression is *ab initio* repressed by thiamine (*thiAp-sec24*). Notice the non-cortical fluorescent puncta, as well as, the total lack of PM-associated signal, indicating a block in UapA and ChsB exocytosis at the ER-exit sites. (D) Epifluorescence microscopy co-localization analysis and relevant quantification (PCC=0.61, P<0.0001, n=5) of a strain co-expressing Sec24-mRFP and UapA-GFP, under ongoing exocytic conditions *(alcAp-uapA,* 90 min derepression). Notice that the partially functional mRFP-tagged version of Sec24 (evident by the growth test shown in A) elicits an apparent delay in UapA exocytosis, reflected in the appearance of prominent labeling of uniform perinuclear ER.

### UapA traffic to the PM by a novel route

We have previously reported that UapA sorting to the PM is independent of AP-1 and AP-3complexes, and clathrin light chain ClaL, which are all essential for post-Golgi traffic of several other cargoes, but depends on clathrin heavy chain (ClaH) *(18).* The involvement of ClaH was difficult to rationalize, because the major function of this protein is the formation of AP-1-coated vesicles from the TGN, the endosome or the PM. Thus, here, we examined whether *ab initio* establishment of repression of ClaH has an effect on the localization specifically of *de novo* synthesized UapA. Additionally, we also followed the effect of *ab initio* repression of RabE^Rab11^ and AP-1 under exactly the same conditions. The logic of testing RabE is that it is known to function as an essential upstream TGN effector essential for the formation of AP-1/clathrin vesicles and post-Golgi cargo secretion.

**Figure 4A, B** shows that repression of AP-1 and RabE did not abolish UapA translocation to the PM after its transcriptional derepression. Contrastingly, in the absence of ClaH, UapA remained within a cytosolic mesh even after prolonged expression (12-14 h). This result confirms that while ClaH is critical for the trafficking of *de novo* made UapA to the PM, two TGN-localized upstream effectors of ClaH function, AP-1 and RabE, proved redundant for UapA secretion. In contrast to the redundancy of RabE and AP-1 but essentiality of ClaH for UapA secretion, the polar localization of a standard apical cargo (e.g. SynA) was abolished in the absence of all three Golgi/TGN-related proteins (**Fig. 4C**), in line with previous reports *(15, 24).*

**Fig. 4.**
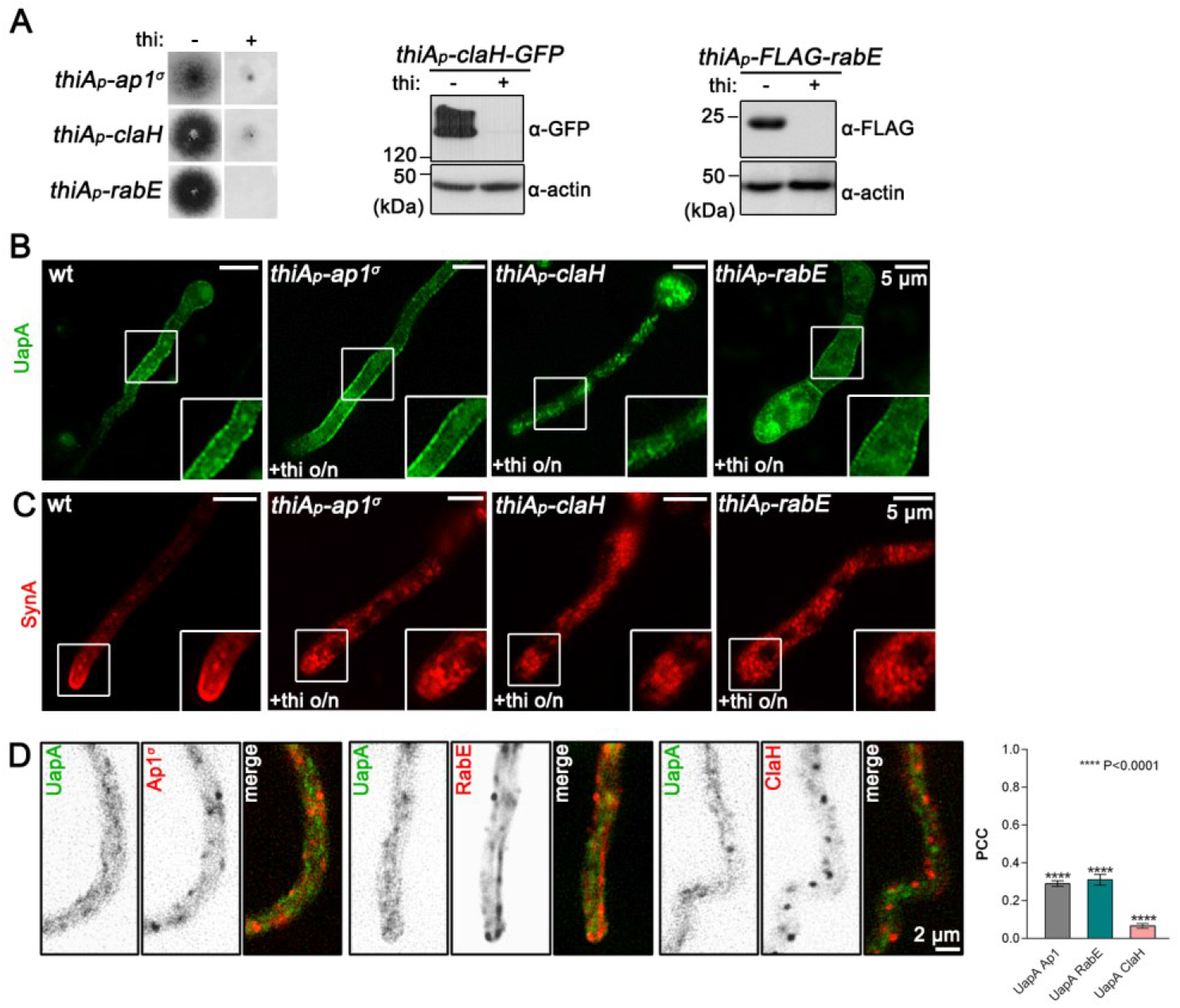
UapA translocation to the PM takes place without involvement of a conventional post-Golgi machinery. (A) Left panel: Growth of isogenic strains expressing UapA-GFP in a background where the Ap1^σ^, ClaH or RabE are expressed under the control of the thiamine regulated promoter *(thiAp-ap1^σ^, thiAp-claH, thiAp-rabE)*, in the absence (-) or presence (+) of thiamine. Middle and Right panels: Western blot analysis comparing protein steady state levels of GFP-or FLAG-tagged versions of ClaH and RabE, respectively, in the absence (-) or presence (+) of thiamine, added *ab initio* for 18 h (overnight culture). Equal protein loading is indicated by actin levels. (B) Epifluorescence microscopy analysis examining the subcellular localization of *de novo* made UapA-GFP in genetic backgrounds lacking AP-1, ClaH and RabE *(thiAp-ap1^σ^, thiAp-claH, thiAp-rabE)* due to pre-established transcriptional repression of the relevant genes (*thiAp-ap1^σ^, thiAp-claH, thiAp-rabE*) by thiamine. (C) Epifluorescence microscopy following the subcellular localization of mCherry-SynA in genetic backgrounds repressed for Ap1^σ^, ClaH and RabE *(thiAp-ap1^σ^, thiAp-claH, thiAp-rabE).* (D) Epifluorescence microscopy co-localization analysis and relevant quantification of strains co-expressing differentially tagged fluorescent versions of UapA and AP-1, RabE or ClaH, under conditions of ongoing UapA exocytosis (*alcAp*, 90 min derepression). For co-localization with AP-1/RabE/ClaH: PCC=0.29/0.31/0.07, P<0.0001, n=7.

We also performed co-localization studies of *de novo* made UapA with RabE, AP-1 and ClaH (**Fig. 4D**). Statistical evaluation of the images obtained showed that there is no co-localization of UapA with AP-1 and RabE (PCC < 0.3) or ClaH (PCC < 0.1). While the non-colocalization of UapA with AP-1 or RabE might have been expected given the redundancy of these Golgi proteins for UapA secretion, the very clear non-colocalization of UapA and ClaH, despite the critical role of the latter in UapA secretion, strongly suggested that ClaH plays an indirect role in secretion, other than cargo vesicle coat packaging at the TGN, as also shown in mammals *(25, 26).*

### UapA sorting to the PM is microtubule independent, but actin-dependent

Conventional cargo secretion in *A. nidulans* and other eukaryotes is known to be microtubule and actin network dependent *(27, 28).* To investigate whether UapA secretion is dependent on distinct components of the cytoskeleton, we used drugs that block microtubule (benomyl) or actin (latrunculin B) organization. **Figure 5A** shows that addition of benomyl, which had destructing effect on tubulin polymerization, had absolutely no effect on *de novo* made UapA translocation to the PM, after 190 min of derepression. On the other hand, benomyl inhibited, as might have been expected the apical localization of a standard apical cargo, such as ChsB (**Fig. 5B**). Contrastingly, addition of latrunculin B led to severe block in the proper secretion of both UapA and ChsB (**Fig. 5C** and **Fig. 5D**, respectively). Thus, proper actin network organization, but not microtubule organization, is required for UapA trafficking to the PM.

**Fig. 5.**
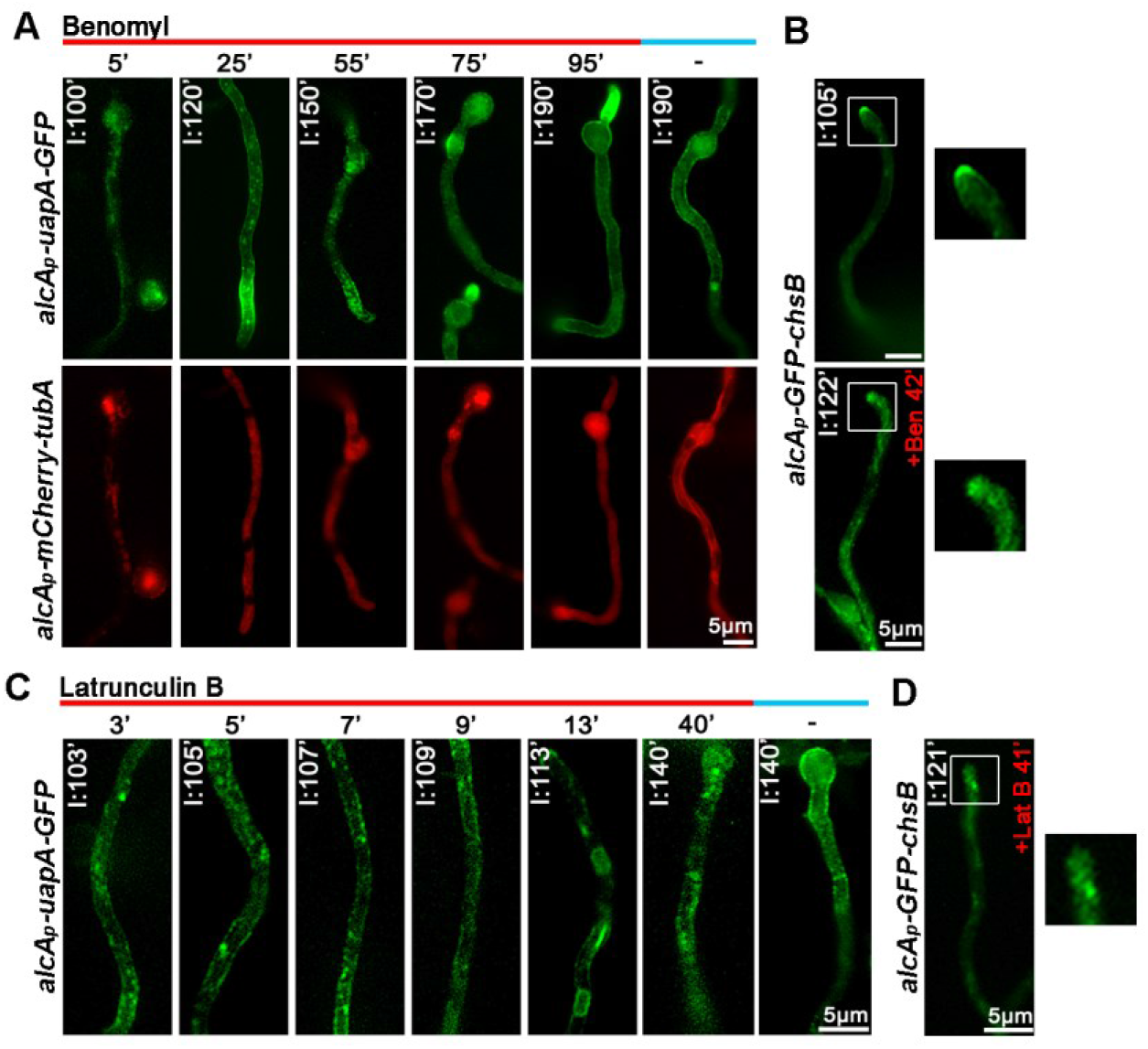
Actin organization, but not microtubules, are essential for UapA translocation in the PM. (A) Time course of treatment of strains expressing UapA-GFP and mCherry-TubA, with the anti-microtubule drug benomyl for 5, 25, 55, 75 and 95 minutes. Localization of UapA followed after its derepression, for a total period of 100, 120, 150, 170 or 190 minutes. In all cases benomyl was added at 95 min of UapA derepression. Notice the appearance of exocytic fluorescent puncta at 100 min, 5 min after benomyl addition. UapA reaches normally the PM despite the absence of functional microtubules, evident by the diffuse cytoplasmic signal of mCherry-TubA (lower panels). An untreated strain is included as control (190 min of derepression, no benomyl addition). (B) Subcellular localization of *alcAp-GFP-chsB* in the absence (upper panel) or presence (42 min, lower panel) of benomyl, after 105 or 122 minutes of induction, respectively. Notice an apparent block of proper localization of ChsB at the hyphal tip. (C) Time course of treatment with the antiactin drug latrunculin B for 3, 5, 7, 9, 13 and 40 minutes, of a strain expressing UapA-GFP under conditions of derepression for 103, 105, 107, 109, 113 or 140 minutes, compared to an untreated strain included as control (140 min). In all cases latrunculin B is added at 100 min of UapA derepression. Notice an apparent block in UapA sorting to the PM in the presence of latrunculin B. (D) Epifluorescence microscopy localization of *alcAp-GFP-chsB* in the presence (41 min) of latrunculin B, after 121 minutes of derepression. Notice the abnormal distribution of ChsB at the hyphal tip, compared to the wt shown in 5B.

### Endosomes are redundant for UapA trafficking to the PM

The non-dependence of UapA secretion on microtubule organization prompted us to examine whether early endosomes (EE) or recycling endosomes (RE), which are key carriers of microtubule-dependent apical cargo sorting and recycling, play any role in the trafficking of *de novo* made UapA. The key Rab GTPase for endosome functioning in cargo sorting in the PM is Rab5. *A. nidulans* has two Rab5 paralogues, RabA and RabB, which co-localize to EEs *(19).* We followed the subcellular localization of *de novo* made UapA in strains that carry a total deletion of *rabB (ΔrabB),* or additionally to *ΔrabB* also carry a thiamine-repressible *rabA* allele (*thiAp-rabA).* **Figure 6A** shows that in both strains UapA is properly localized to the PM. In line with this, transient UapA-labeled cytosolic structures, observed soon after the *de novo* synthesis of UapA, did not co-localize with RabB-marked endosomes (**Fig. 6B**). Overall, these results confirmed that trafficking of newly made UapA is independent of early/recycling endosome functioning.

**Fig. 6.**
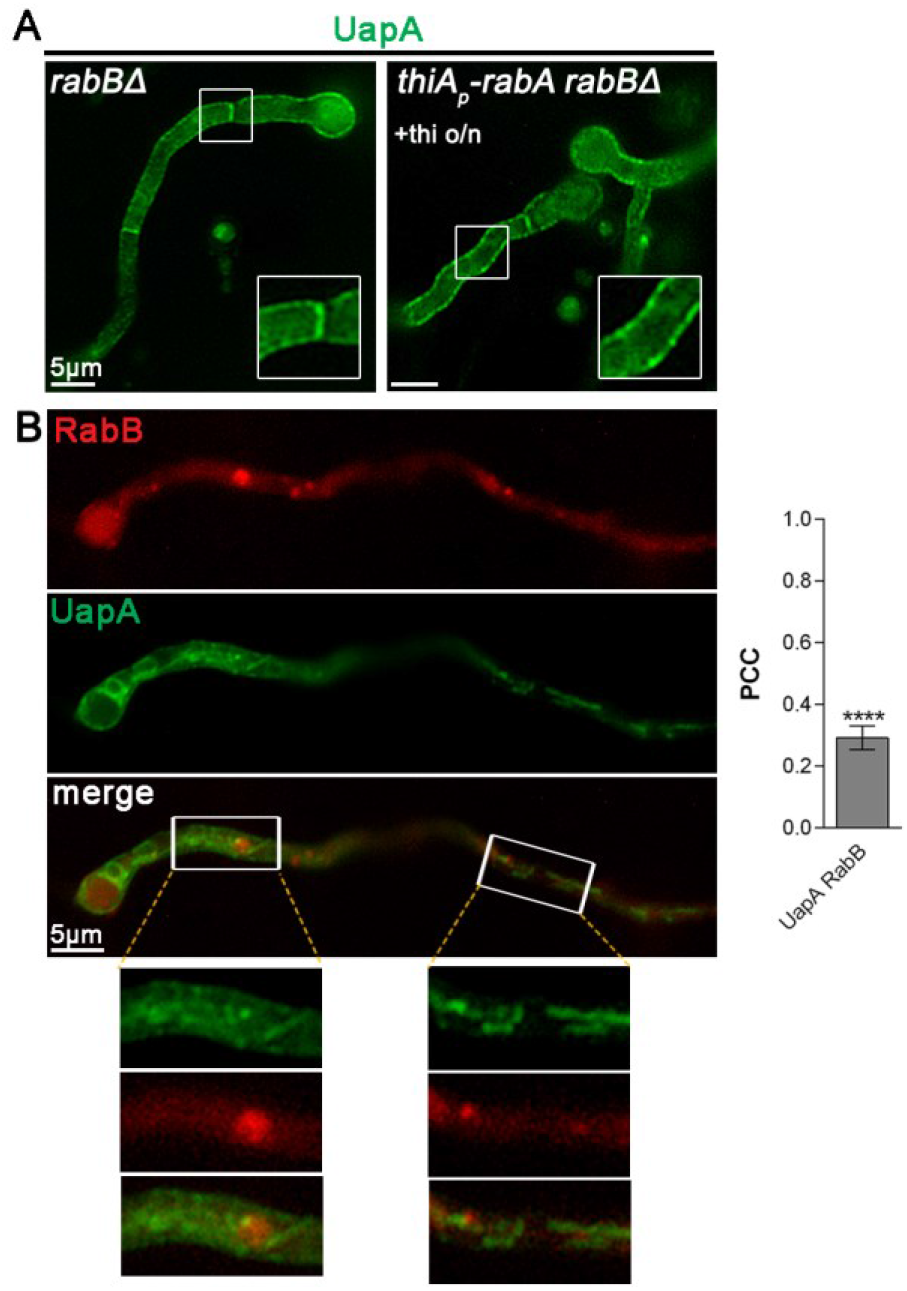
Early endosomes are dispensable for UapA exocytosis. (A) Epifluorescence microscopy analysis of UapA-GFP localization in the absence of either only RabB (left panel) or both RabA and RabB (right panel). RabA is *ab initio* repressed (o/n) via the thiamine regulated promoter *thiAp.* Notice that the transporter is sorted to the PM normally. (B) Relative subcellular localization and quantification (PCC=0.29, P<0.0001, n=8) of RabB-mRFP and UapA-GFP co-localization, under exocytic conditions *(alcAp-uapA,* 90 min derepression). Notice the absence of significant colocalization between the two proteins, highlighting the non-endosomal character of UapA exocytic vesicles.

## Discussion

Here we uncover a novel trafficking route concerning nutrient transporter translocation to the PM that proved independent of a) Golgi/TGN functioning, b) acquisition of post-Golgi secretory vesicle identity (i.e. independent of Rab11 and AP-1/clathrin coating), c) microtubule organization, and d) endosomes. This route requires functional ERes (i.e. COPII-dependent) and actin network organization. Additionally, it requires a function of ClaH that is not related to Rab11/AP-1-dependent coated vesicle formation the TGN, but is rather connected with proper actin organization, as reported in mammalian cells *(25, 26).* The above findings are in line with the observation that in the course of its secretion, *de novo* made UapA transporter is never detected in Golgi and/or endosomal compartments. Following this Golgi-independent secretory route, UapA and seemingly other nutrient transporters (**Fig. S2**) are localized, rather homogeneously, all along the hyphal cell PM. In other words, unlike apical membrane cargoes, which show polar secretion towards actively growing hyphal tips, transporters do not show any indication of polarization in their PM localization.

Alternative pathways of membrane secretion have also been described for a limited number of specific cargoes, but these unconventional routes are still Golgi/TGN-dependent *(11, 29–31).* The best-documented example of a protein sorting that bypasses the Golgi is the CTFR transporter, which is associated with cystic fibrosis, one of the most common lethal genetic disorders in humans. CFTR translocates to the PM via direct trafficking from the ER in COPII vesicles *(11–13).* This pathway, similar to findings presented herein, does not depend on Sed5, Arf1, or Rab, effectors essential for anterograde transport in the early secretory pathway. Another example of a cargo that is secreted in Golgi-independent, but also COPII-independent, manner is the yeast transmembrane protein Ist2, an essential protein of the cortical ER and an ER-to-PM tethering factor *(11).* Interestingly, Ist2 seems to be required for efficient trafficking from the ER to the PM of newly synthesized leucine transporter Bap2 *(32).* Some signal-peptide-containing proteins also reach the cell surface in a COPII- and Golgi-independent manner *(11).* Noticeably, the pivotal role of actin dynamics in transporter PM sorting that is evidenced here is similar to what has been observed for a number of other transporters, such as the GLUT4 glucose transporter *(33, 34).*

A non-polar route taking care of transporter trafficking to the PM has a strong rational basis. Most transporters are localized to the PM in order to feed cells with nutrients from their environment. Thus, there is no obvious need to drive transporters to a specialized part of the PM, such as the actively growing tip of fungi, where other specific proteins involved in PM and cell wall biosynthesis are localized via the conventional polarized secretion route. By analogy to filamentous fungi, in other more complex cells that also face the challenge of polar growth, as for example neurons, transporters involved in synaptic neurotransmission are expected to be polarly localized via conventional routes in the synaptic region, but transporters serving specifically nutrient supply and dendrite or soma homeostasis might well be sorted non-polarly, via a mechanism similar to the one we discovered here *(35, 36).* Notably also, in dendrites, forward secretory trafficking of specific membrane cargoes (e.g. glutamate receptor GluA1 and neuroligin 1) seems to occur via a Golgi-independent route, which however involves a type of sorting endosomes *(37).*

Evidently, several questions concerning the mechanism of transporter trafficking revealed by our findings remain to be addressed. What makes ER-budding vesicles carrying transporters distinct from conventional COPII vesicles carrying polar cargos? Are transporter vesicles loaded to actin filaments coated or uncoated? How clathrin and actin remodeling might orchestrate interactions around the transporter vesicle as its approaches the PM? A key logical assumption for answering some of the above questions would be to consider that the transporters themselves contain intrinsic information for their particular trafficking mechanism. Transporters are large polytopic transmembrane proteins, which often partition into specific lipid domains or oligomerize early after their translocation in the ER membrane. Importantly, transporter oligomerization, including that of UapA, has been shown to be critical for ER-exit *(38, 39).* This is in line with results supporting that transmembrane protein oligomerization can mechanistically drive membrane curvature and vesicular budding, even in the absence of vesicular coats *(40).* We predict that the intrinsic capacity of transporter oligomerization might be necessary and sufficient for ER-exit and formation of specific transporter vesicles. Once a transporter vesicle is properly formed, no other control point exists to stop its bulk trafficking to the actin network, and from there to the PM. This nicely explains why partial misfolding and improper oligomerization of transporters lead to ER-retention, but not to retention in any other subcellular compartment. In some cases, as in the case of yeast amino acid transporters, chaperone-assisted oligomerization might also needed for proper ER exit *(41).*

## Acknowledgments

We are grateful to the Erasmus student Valentina D’ Autilia for help in experiments concerning transporters other than UapA. We also thank R. Fischer for providing the myosin knock-out strains and to S. Efthmiopoulos for microscopy facilities.

## Funding

The current research was supported by the *Fondation Santé,* to which we are grateful.

## Author contributions

V. Bouris: Investigation, Visualization. O. Martzoukou: Validation, Formal Analysis, Investigation, Visualization. S. Amillis: Investigation, Visualization. G. Diallinas: Conceptualization, Writing – original draft preparation, Writing – review and editing, Visualization, Supervision, Project administration, Funding acquisition, Conceptualization.

## Competing interests

Authors declare no competing interests.

## Data and materials availability

All data is available in the main text or the supplementary materials.

## List of Supplementary Materials

Materials and Methods Supplementary text Figs. S1-S2 Tables S1-S2 Movies S1-S2

