## Supplementary material for "Nutrient transporter translocation to the plasma membrane via a Golgi-independent unconventional route"

Science
AAAS

### Supplementary Materials for

#### Nutrient transporter translocation to the plasma membrane via a Golgi-independent unconventional route

**Authors:** Vangelis Bouris[1#], Olga Martzoukou[1#], Sotiris Amillis[1] and George Diallinas[1*].

**Affiliations**:

[1]Department of Biology, National and Kapodistrian University of Athens, Panepistimioupolis 15784, Athens, Greece.

[#]Equal contribution

**This PDF file includes:**

Materials and Methods
Supplementary text
Figs.S1 to S2
Tables S1 to S2
Captions for Movies S1 to S2

**Other Supplementary Materials for this manuscript include the following:**

Movies S1 to S2

#### Materials and Methods

Media, strains, growth conditions and transformation

Standard complete and minimal media for *A. nidulans* were used (FGSC, http://www.fgsc.net.). Media and chemical reagents were obtained from Sigma-Aldrich (Life Science Chemilab SA, Hellas) or AppliChem (Bioline Scientific SA, Hellas). Glucose 0.1-1 % (w/v) or Fructose 0.1 % (w/v) was used as carbon sources. $NH_4^+$ (di-ammonium tartrate) and $NaNO_3$ were used as nitrogen sources at 10 mM. Thiamine hydrochloride was used at a final concentration of 10 μM as a repressor of the *thiA*$_p$ promoter. *A. nidulans* transformation was performed by generating protoplasts from germinating conidiospores as described previously in Koukaki *et al. (42)*, using TNO2A7 as a recipient strainthat allow selection of transformants via complementation of a pyrimidine autotrophy *(43)*. Integrations of gene fusions with fluorescent tags, promoter replacement fusions, or deletion cassettes were selected using the *A. fumigatus* markers orotidine-5'-phosphate-decarboxylase (*AFpyrG*, Afu2g0836), GTP-cyclohydrolase II (*AFriboB*, Afu1g13300) or a pyridoxine biosynthesis gene (*AFpyroA*, Afu5g08090), resulting in complementation of the relevant auxotrophies. Transformants were verified by PCR and Southern analysis. Combinations of mutations and fluorescent epitope-tagged strains were generated by standard genetic crossing and progeny analysis. *E. coli* strains used were dH5α. *A. nidulans* strains used are listed in **Table S1**.

Nucleic acid manipulations and plasmid constructions

Genomic DNA extraction was performed as described in FGSC (http://www.fgsc.net). All DNA fragments used in the various constructs were amplified from a TNO2A7 strain. Plasmid preparation and DNA gel extraction were performed using the Nucleospin Plasmid and the Nucleospin Extract II kits (Macherey-Nagel, Lab Supplies Scientific SA, Hellas). Restriction enzymes were from Takara Bio or Minotech (Lab Supplies Scientific SA, Hellas). DNA sequences were determined by Eurofins-Genomics (Vienna, Austria). Conventional PCR reactions and high fidelity amplifications were performed using KAPA Taq DNA and Kapa HiFi polymerases (Kapa Biosystems, Roche Diagnostics, Hellas). Gene cassettes were generated by sequential cloning of the relevant fragments in the pGEM-T plasmid, which served as template to PCR-amplify the relevant linear cassettes. For primers see **Table S2**.

Protein extraction and western blots

Total protein extraction was performed as previously described using dry mycelia from cultures grown in minimal media supplemented with $NaNO_3$ at 25° C *(44)*. Total proteins (50 μg, estimated by Bradford assays) were separated in a 10 % (w/v) polyacrylamide gel and were transferred on PVDF membranes (GE Healthcare Life Sciences Amersham). Immunodetection was performed with an anti-FLAG M2 monoclonal antibody (Sigma-Aldrich), an anti-GFP monoclonal antibody (Roche Diagnostics), an anti-actin monoclonal (C4) antibody (MP Biomedicals Europe) and an HRP-linked antibody (Cell Signaling Technology Inc). Blots were developed using the LumiSensor Chemiluminescent HRP Substrate kit (Genscript USA) and SuperRX Fuji medical X-Ray films (FujiFILM Europe).

Fluorescence Microscopy and Statistical Analysis

Samples were prepared as previously described *(15)*. Unless otherwise stated, conidiospores were incubated overnight in glass bottom 35mm μ-dishes (*ibidi*, Lab Supplies Scientific SA, Hellas) in liquid minimal media, for 16-22 h at 25° C, under conditions of transcriptional repression of *uapA* (10 mM $NH_4^+$ when expressed from its native promoter or 1% glucose when expressed from $alcA_p$-*uapA*) and of genes involved in trafficking expressed under the $thiA_p$ promoter (10 μM thiamine). Transcriptional repression of *uapA* was followed by a derepression period, through a shift in media containing either $NaNO_3$ as a nitrogen source for native *uapA,* or fructose as a carbon source for $alcA_p$-*uapA*. Derepression periods ranged from to 30 min to 12 h, according to experiments. Benomyl (Sigma-Aldrich), Brefeldin A (Cayman Chemical) and Latrunculin B (Sigma-Aldrich) were used at 2.5μg $ml^{-1}$, 100μg $ml^{-1}$ and 100μg $ml^{-1}$, final concentrations respectively. FM4-64 (Thermo Fischer Scientific) staining was according to Martzoukou *et al. (18)*.Images were obtained using an inverted Zeiss Axio Observer Z1 equipped with an Axio Cam HR R3 camera. Contrast adjustment, area selection and color combining were made using the Zen lite 2012 software. Scale bars were added using the FigureJ plugin of the ImageJ software. Images were further processed and annotated in Adobe Photoshop CS4 Extended version 11.0.2. For quantifying colocalization, Pearson's correlation coefficient (PCC) above thresholds, for a selected Region of interest (ROI) was calculated using the ICY co-localization studio plugin (pixel-based method) (http://icy.bioimageanalysis.org/). One sample t-test was performed to test the significance of differences in PCCs, using the Graphpad Prism software. Confidence interval was set to 95%.

#### Supplementary text

Several transporters follow the unconventional secretion pathway of UapA

Using UapA as a model cargo we uncovered a novel sorting pathway concerning a nutrient transporter. We wanted to investigate whether our findings also hold true for other PM transporters. Thus, we selected to study the sorting of a number of structurally, functionally and evolutionary distinct transporters. These include the AzgA purine transporter *(45)*, the FcyB cytosine-purine transporter (*46)* the FurA allantoin transporter *(47)* and the L-proline PrnB transporter *(48)*. Homologues of these transporters are ubiquitously conserved in most domains of life. **Fig. S2** highlights the results obtained after following the sorting of these transporters in the presence of benomyl, latrunculin B, or when AP-1 or RabE are transcriptionally knocked-down. All transporters tested were properly sorted in the PM in the absence of AP-1 and RabE, or in the presence of benomyl, but not when latrunculin B disorganizes actin. These findings strongly suggest that nutrient transporter trafficking to the PM occurs by a Golgi-independent, non-polar, mechanism identical to that discover with of UapA.

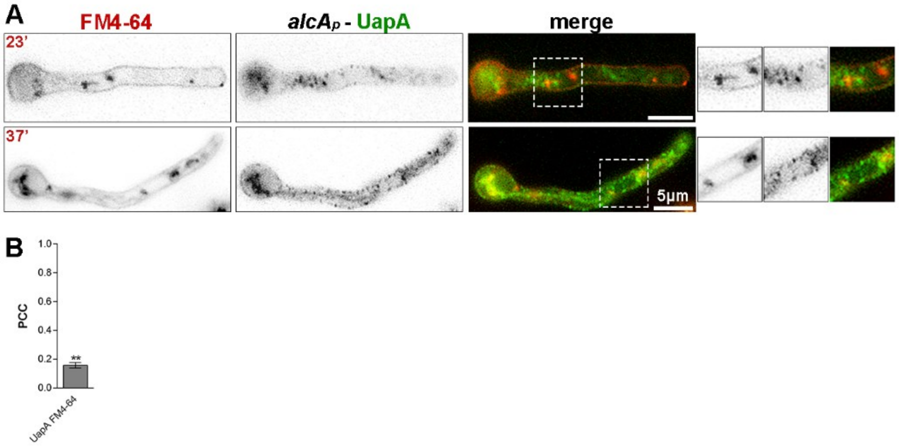


**Fig. S1.**
(A) Subcellular localization of *de novo* made UapA-GFP (*alcA*$_p$-*uapA*, 90 min derepression) in the presence of the lipophilic endocytic dye FM4-64 for 23 and 37 min (upper and lower panels, respectively). (B) Relevant quantification of co-localization (PCC=0.16, P=0.0036, n=4). Notice that UapA-labeled exocytic vesicles do not co-localize significantly with FM4-64.

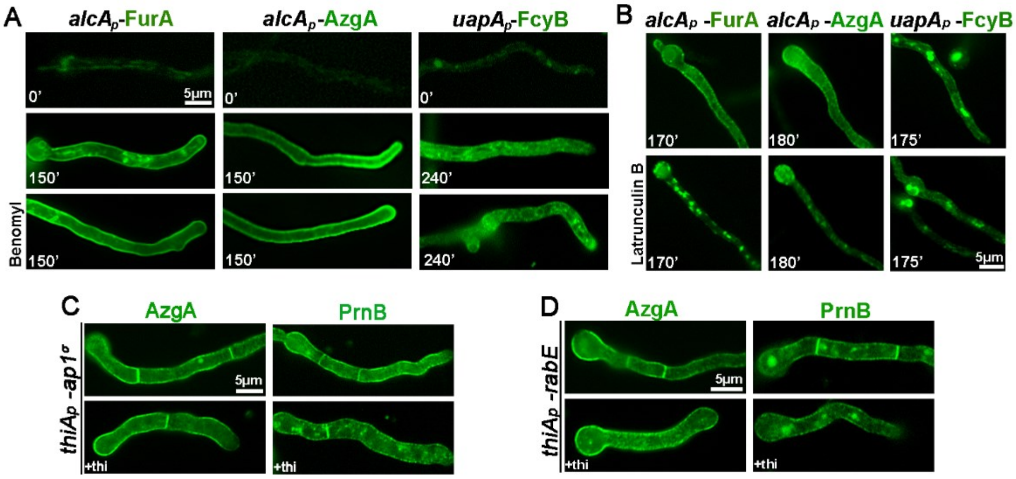


**Fig. S2.**

(A, B) Epifluorescence microscopy analysis of the exocytosis of three GFP-tagged transporters, namely FurA, AzgA and FcyB (see text), in the absence or presence of Benomyl (A) or Latrunculin B (B), after derepression of the relevant genes for the time indicated. Notice that addition of Benomyl has no effect on UapA translocation to the PM, whereas, Latrunculin B has a significant negative effect on UapA secretion. (C, D) Epifluorescence microscopy images showing that sorting of AzgA or PrnB to the PM does not require two basic components of the post-Golgi exocytic machinery, AP-1 (C) and RabE (D), respectively.

**Table S1.**

Strains used in this study. All strains carry the *veA1* mutation affecting sporulation. *pabaA1*, *pyroA4*, *riboB2*, *argB2*, *pyrG89, pantoB100, biA1, nicA2*and *inoB2* are auxotrophic mutations for p-aminobenzoic acid, pyridoxine, riboflavin, arginine, uracil/uridine, D-pantothenic acid, biotin, nicotinic acid and inositol respectively. *yA2* and *wA3* or *wA4*are mutations resulting in yellow and white conidiospore colors respectively.

| **Name** | **Genotype** | **Reference** |
|---|---|---|
| TNO2A7 | nkuAΔ::argBpyrG89 pyroA4 riboB2 | 43 |
| Δ7 | uapAΔ uapCΔ::AFpyrG azgAΔ fcyBΔ::argB furDΔ::AFriboB furAΔ::AFriboB cntAΔ::AFriboB pantoB100 pabaA1 | 49 |
| mRFP-PH$^{OSBP}$ | pyroA4[pyroA::gpdA$^{m}{}_{p}$::mRFP-PH$^{OSBP}$] inoB2 niiA4 wA4 | 23 |
| mCherry-sedV | pyroA4[pyroA::gpdA$^{m}{}_{p}$::mcherry-sedV] nkuAΔ::bar, wA4, niiA4 inoB2 | 20 |
| mCherry-synA | AFpyG-mcherry-synA yA::AFpyroA GFP-tpmA fwA1 pyrG89 pyroA4 nicA2 nkuAΔ::argB | 24 |
| sedVts | sedVR238G::AFpyrG pyroA4 pyrG89 nkuAΔ::bar | 22 |
| mCherry-tubA | alcA$_p$::mCherry-tubA::pyroA nkuAΔ::argB pyrG89 pyroA4 | 50 |
| alcA$_p$-mRFP-rabB | alcA$_p$-mRFP-rabB::pyroA nkuAΔ::bar inoB2 pyroA4 niiA4 wA4 | 51 |
| uapA-GFP | uapAΔ::uapA-GFP::AFriboB uapCΔ::AfpyrG nkuAΔ::argB pabaA1 pyroA4 riboB2 | 10 |
| alcA$_p$-uapA-GFP | uapAΔ::alcA$_p$::uapA-GFP::AFriboB uapCΔ::AfpyrG nkuAΔ::argB pabaA1 pyroA4 riboB2 | 10 |
| fcyB-GFP | (pBS-argB)fcyB-GFP uapAΔ uapCΔ::AFpyrG azgAΔ argB2 pabaA1 | 46 |
| prnB-GFP | prnBΔ::prnB-GFP argB2 pabaA1 yA2 | 52 |
| alcA$_p$-GFP-chsB | alcAp-GFP-chsB::Ncpyr4 nkuAΔ::argB pyrG89 pyroA4 | 50 |
| alcA$_p$-uapA-GFP artAΔ | uapAΔ::alcA$_p$-uapA-GFP::AFriboB artAΔ::AFriboB nkuAΔ::argB pyroA4 | This study |
| GFP-chsB | GFP-chsB::AFpyrG nkuAΔ::argB pyrG89 pyroA4 riboB2 | This study |
| alcA$_p$-uapA-GFP sedV$^{ts}$ | (pBS-argB)-alcAp-uapA-GFP sedVR238G::AFpyrG uapAΔ pabaA1 pyroA4 | This study |
| alcA$_p$-furA-GFP | (pGEM-alcA$_p$-panB)alcA$_p$-furA-GFP uapAΔ uapCΔ::AFpyrG azgAΔ fcyBΔ::argB furDΔ::AFriboB furAΔ::AFriboB cntAΔ::AFriboB pantoB100 pabaA1 | This study |
| alcA$_p$-azgA-GFP | (pGEM-alcA$_p$-panB)alcA$_p$-azgA-GFP uapAΔ uapCΔ::AFpyrG azgAΔ fcyBΔ::argB furDΔ::AFriboB furAΔ::AFriboB cntAΔ::AFriboB pantoB100 pabaA1 | This study |
| alcA$_p$-uapA-GFP mRFP-PH$^{OSBP}$ | pyroA::gpdA$^{m}{}_{p}$::mRFP-PH$^{OSBP}$uapAΔ::alcA$_p$-uapA-GFP::AFriboB pabaA1 inoB2 | This study |
| alcA$_p$-uapA-GFP mCherry-sedV | pyroA::gpdA$^{m}{}_{p}$::mcherry-sedV uapAΔ::alcA$_p$-uapA-GFP::AFriboB pabaA1 inoB2 | This study |
| alcA$_p$-uapA-GFP sec24-mRFP | sec24-$^{(5xGA)}$mRFP::AFpyrG pyrG89 uapAΔ::alcA$_p$-uapA-GFP::AFriboB nkuAΔ::argB pabaA1 pyroA4 riboB2 | This study |
| alcA$_p$-uapA-GFP ap1$^{\sigma}$-mRFP | ap1$^{\sigma}$-$^{(5xGA)}$mRFP::AFpyrG uapAΔ::alcA$_p$-uapA-GFP::AFriboB nkuAΔ::argB pyrG89 pyroA4 riboB2 | This study |
| alcA$_p$-uapA-GFP mRFP-rabE | mRFP-rabE::AFpyrG uapAΔ::alcA$_p$-uapA-GFP::AFriboB nkuAΔ::argB pyrG89 pyroA4 | This study |

| | | |
|---|---|---|
| | riboB2 | |
| alcA$_p$-uapA-GFP claH-mRFP | claH-$^{(5xGA)}$mRFP::AFpyrG uapAΔ::alcA$_p$-uapA-GFP::AFriboB nkuAΔ::argB pyrG89 pyroA4 riboB2 | This study |
| alcA$_p$-uapA-GFP mCherry-tubA | alcA$_p$-mCherry-tubA::pyroA uapAΔ::alcA$_p$-UapA-GFP::AFriboB nkuAΔ::argB pabaA1 | This study |
| alcA$_p$-uapA-GFP alcA$_p$-mRFP-rabB | alcA$_p$-mRFP-rabB::pyroA uapAΔ::alcA$_p$-uapA-GFP::AFriboB wA3 pabaA1 | This study |
| thiA$_p$-sedV uapA-GFP | thiA$_p$-sedV::AFpyrG uapAΔ::uapA-GFP nkuAΔ::argB pyrG89 pyroA4 | This study |
| thiA$_p$-hypB uapA-GFP | thiA$_p$-hypB::AFpyrG uapAΔ::uapA-GFP nkuAΔ::argB pyrG89 pyroA4 | This study |
| thiA$_p$-geaA uapA-GFP | thiA$_p$-geaA::AFpyrG uapAΔ::uapA-GFP nkuAΔ::argB pyrG89 pyroA4 | This study |
| thiA$_p$-rabC uapA-GFP | thiA$_p$-rabC::AFpyrG uapAΔ::uapA-GFP nkuAΔ::argB pyrG89 pyroA4 | 18 |
| thiA$_p$-rabO uapA-GFP | thiA$_p$-rabO::AFpyrG uapAΔ::uapA-GFP nkuAΔ::argB pyrG89 pyroA4 | This study |
| thiA$_p$-sec24 alcA$_p$-uapA-GFP | thiA$_p$-sec24::AFpyrG uapAΔ::alcA$_p$-uapA-GFP nkuAΔ::argB pyrG89 pyroA4 | This study |
| thiA$_p$-sec24 GFP-chsB | thiAp-sec24::AFriboB GFP-chsB::AFpyrG nkuAΔ::argB pyrG89 pyroA4 riboB2 | This study |
| thiA$_p$-sedV GFP-chsB | thiAp-sedV::AFriboB GFP-chsB::AFpyrG nkuAΔ::argB pyrG89 pyroA4 riboB2 | This study |
| thiA$_p$-hypB GFP-chsB | thiAp-hypB::AFriboB GFP-chsB::AFpyrG nkuAΔ::argB pyrG89 pyroA4 riboB2 | This study |
| thiA$_p$-geaA GFP-chsB | thiAp-geaA::AFriboB GFP-chsB::AFpyrG nkuAΔ::argB pyrG89 pyroA4 riboB2 | This study |
| thiA$_p$-rabC GFP-chsB | thiAp-rabC::AFriboB GFP-chsB::AFpyrG nkuAΔ::argB pyrG89 pyroA4 riboB2 | This study |
| thiA$_p$-rabO GFP-chsB | thiAp-RabO::AFriboB GFP-chsB::AFpyrG nkuAΔ::argB pyrG89 pyroA4 riboB2 | This study |
| thiA$_p$-ap1$^{\sigma}$alcA$_p$-uapA-GFP | thiA$_p$::$^{FLAG}$ap1$^{\sigma}$::AFriboB (pBS-argB)alcA$_p$-uapA-GFP pabaA1 | 15 |
| thiA$_p$-claH uapA-GFP | uapAΔ::uapA-GFP::AFriboB thiA$_p$-claH::AFpyroA nkuAΔ::argB pyroA4 pabaA1 | 15 |
| thiA$_p$-rabE uapA-GFP | thiA$_p$-rabE::AFpyrG uapAΔ::uapA-GFP nkuAΔ::argB pyrG89 pyroA4 | This study |
| chs5Δ uapA-GFP | chs5Δ::AFpyrGuapAΔ::uapA-GFPnkuAΔ::argBpyrG89 pyroA4 | This study |
| ggAΔ uapA-GFP | ggAΔ::AFpyrG uapAΔ::uapA-GFP nkuAΔ::argB pyrG89 pyroA4 | This study |
| thiA$_p$-ap1$^{\sigma}$ mCherry-synA | thiAp-ap1σ::AFriboB yA::AFpyroA tpmAp-gfp-tpmA AFpyG::mcherry-synA nkuAΔ::argB pyrG89 | This study |
| thiA$_p$-claH mCherry-synA | ap2σ-(5xGA)GFP::AFpyrG claH::thiAp-claH::AFpyroA AFpyG::mcherry-synAnkuAΔ:argB pyrG89 pyroA4 | This study |
| thiA$_p$-rabE mCherry-synA | thiAp-rabE::AFriboB AFpyrG::mcherry-synA nkuAΔ::argB pyrG89 pyroA4 | This study |
| alcA$_p$-uapA-GFP rabBΔ thiA$_p$-rabA | thiA$_p$-rabA::AFpyroA rabBΔ::AFpyrG uapAΔ::alcA$_p$-uapA-GFP::AFriboB nkuAΔ::argB pyrG89 pyroA4 riboB2 | This study |
| thiA$_p$-FLAG-hypB | thiA$_p$-$^{FLAG}$-hypB::AFpyrG uapAΔ::uapA-GFP nkuAΔ::argB pyrG89 pyroA4 | This study |
| thiA$_p$-FLAG-geaA | thiA$_p$-$^{FLAG}$-geaA::AFpyrG uapAΔ::uapA-GFP nkuAΔ::argB pyrG89 pyroA4 | This study |
| thiA$_p$-FLAG-rabC | thiA$_p$-$^{FLAG}$-rabC::AFpyrG uapAΔ::uapA-GFP nkuAΔ::argB pyrG89 pyroA4 | This study |
| thiA$_p$-FLAG-rabE | thiA$_p$-$^{FLAG}$-rabE::AFpyrG uapAΔ::uapA-GFP nkuAΔ::argB pyrG89 pyroA4 | This study |
| thiA$_p$-FLAG-rabO | thiA$_p$-$^{FLAG}$-rabO::AFpyrG uapAΔ::uapA-GFP nkuAΔ::argB pyrG89 pyroA4 | This study |

| | | |
|---|---|---|
| thiA$_p$-claH-GFP | *AFpyroA::thiAp-claH-(5xGA)GFP::AFpyrG pyrG89 pyroA4 riboB2* | 15 |
| thiA$_p$-ap1$^\sigma$azgA-GFP | *thiA$_p$-FLAG-ap1$^\sigma$::AFriboB azgA-GFP pabaA1 pyroA4* | 18 |
| thiA$_p$-ap1$^\sigma$prnB-GFP | *thiA$_p$-FLAG- ap1$^\sigma$::AFriboB prnB$\Delta$::prnB-GFP pabaA1 pyroA4* | This study |
| thiA$_p$-rabEazgA-GFP | *thiA$_p$-rabE::AFriboB (pGEM-alcA$_p$-panB)alcA$_p$-azgA-GFP azgA$\Delta$* pabaA1 | This study |
| thiA$_p$-rabEprnB-GFP | *thiA$_p$-rabE::AFriboB prnB$\Delta$::prnB-GFP pyrG89* | This study |

**Table S2.**

Oligonucleotides used in this study for cloning purposes.

| Oligonucleotides | Sequence |
|---|---|
| ***pBS SKII claH-$^{(5xGA)}$mRFP::AFpyrG*** | |
| claH ORF KpnI F | CGCGGGTACCCTGGACCAGCTCGCAGAACTTGAAG |
| claH ORF NS SpeI R | CGCGACTAGTGAAAGGACGGAACCCCGTGGCCTG |
| claH 3' SpeI F | CGCGACTAGTGCTCGCCTTGTCTTTTGAGGGGTAG |
| claH 3' NotI R | CGCGGCGGCCGCGGACAATCAGATTGACAGGGAGGG |
| 5xGA SpeI F | CGCGACTAGTGGAGCTGGTGCAGGCGCTGGAGCCGGTGCC |
| AFpyrG SpeI R | CGCGACTAGTACTGTCTGAGAGGAGGCACTGATGCG |
| ***pGEM hypB-$^{(5xGA)}$mRFP::AFpyrG*** | |
| hypB ORF ApaI F | CGCGGGGCCCCTAGGGCCTTGACATACCTTTTCG |
| hypB ORF NS SpeI R | CGCGACTAGTGCGCCGGCCGACGCTATGCTTGCGAG |
| hypB 3' SpeI F | CGCGACTAGTCCGCTTTAGACGGCTCCTATATTCAG |
| hypB 3' NotI R | CGCGGCGGCCGCGAGAAGCGGATGCGAGTTGCCTGAG |
| 5xGA SpeI F | CGCGACTAGTGGAGCTGGTGCAGGCGCTGGAGCCGGTGCC |
| AFpyrG SpeI R | CGCGACTAGTACTGTCTGAGAGGAGGCACTGATGCG |
| ***pGEM GFP-chsB::AFpyrG*** | |
| chsB 5' ApaI F | CGCGGGGCCCGGAGAACAACGACCAGTTGAAATAC |
| chsB 5' SpeI R | CGCGACTAGTGGTTAAACTGGTAGTATGTGCGAATAG |
| chsB ORF SpeI F | CGCGACTAGTATGGCCTACCACGGCTCTGGTC |
| chsB ORF NotI R | CGCGGCGGCCGCGGATTTCACCACAAGCACCTCCG |
| GFP SpeI F | CGCGACTAGTATGGTGAGCAAGGGCGAGG |
| GFPns SpeI R | CGCGACTAGTCTTGTACAGCTCGTCCATG |
| AFpyrG SphI F | CGCGGCATGCGCCTCAAACAATGCTCTTCACCC |
| AFpyrG SphI R | CGCGGCATGCCTGTCTGAGAGGAGGCACTGATG |
| ***pGEM thiA$_p$-sedV::AFpyrG / pGEM thiA$_p$-sedV::AFriboB / pGEM thiA$_p^{FLAG}$-sedV::AFpyrG*** | |
| ***sedV*** 5' ApaI F | CGCGGGGCCCGCCGAGAGACGGATTGTGATGTAAC |
| ***sedV*** 5' SpeI R | CGCGACTAGTGGGTGAATCTAATCGATAAGGGG |
| ***sedV*** ORF SpeI F | CGCGACTAGTATGACCGGGCCTACGATACAGGATC |
| ***sedV*** ORF NotI R | CGCGGCGGCCGCCCTTGCTGAACGCCCGTGTTTAAGG |
| AFpyrG SpeI F | CGCGACTAGTGCCTCAAACAATGCTCTTCACCCTC |
| AFriboB SpeI F | CGCGACTAGTCCCGGGCTGCAGGAATTCG |
| thiA$_p$ SpeI R | CGCGACTAGTGTTGACTCAGTTCAATGGTTCGAC |
| ***pGEM thiA$_p$-hypB::AFpyrG / pGEM thiA$_p$-hypB::AFriboB / pGEM thiA$_p^{FLAG}$-hypB::AFpyrG*** | |
| hypB 5' ApaI F | CGCGGGGCCCGGTTGATGATAGCGGTGAAGCGC |
| hypB 5' SpeI R | CGCGACTAGTCTGTCTCACGAAACAGAACCCCG |
| hypB ORF SpeI F | CGCGACTAGTATGGCGGAAGCTGAGAACGACCC |
| hypB ORF NotI R | CGCGGCGGCCGCCCACATCGTCTTCGTTACTTGAAGC |
| AFpyrG SpeI F | CGCGACTAGTGCCTCAAACAATGCTCTTCACCCTC |
| AFriboB SpeI F | CGCGACTAGTCCCGGGCTGCAGGAATTCG |
| thiA$_p$ SpeI R | CGCGACTAGTGTTGACTCAGTTCAATGGTTCGAC |
| ***pGEM thiA$_p$-geaA::AFpyrG / pGEM thiA$_p$-geaA::AFriboB / pGEM thiA$_p^{FLAG}$-geaA::AFpyrG*** | |
| geaA 5' ApaI F | GTAGAGACGGGCCCGTAGAACAC |
| geaA 5' XbaI R | CGCGTCTAGAGGGCTGTTTACCCGACTGGGATTAG |
| geaA ORF XbaI F | CGCGTCTAGAATGTCTTCCTCCTCTCTGCCAATTG |
| geaA 3' NotI R | CGCGGCGGCCGCGAAGGTCGATGAGCACGCGGAAC |
| AFpyrG SpeI F | CGCGACTAGTGCCTCAAACAATGCTCTTCACCCTC |
| AFriboB SpeI F | CGCGACTAGTCCCGGGCTGCAGGAATTCG |
| thiA$_p$ SpeI R | CGCGACTAGTGTTGACTCAGTTCAATGGTTCGAC |
| ***pGEM thiA$_p$-rabO::AFpyrG / pGEM thiA$_p$-rabO::AFriboB / pGEM thiA$_p^{FLAG}$-rabO::AFpyrG*** | |
| rabO 5' ApaI F | CGCGGGGCCCGAAGGGTTCAAGGATGAAACGAAC |
| rabO 5' XbaI R | CGCGTCTAGACTCAGTGCAGGAGAGCAAAGGAG |
| rabO ORF XbaI F | CGCGTCTAGAATGAACCCTGAGTGGTAAGTGCTTTG |

| | |
|---|---|
| rabO 3' NotI R | CGCGGCGGCCGCCAACCTCTAATTTACCACGGCAGC |
| AFpyrG SpeI F | CGCGACTAGTGCCTCAAACAATGCTCTTCACCCTC |
| AFriboB SpeI F | CGCGACTAGTCCCGGGCTGCAGGAATTCG |
| thiA$_p$ SpeI R | CGCGACTAGTGTTGACTCAGTTCAATGGTTCGAC |
| ***pGEM thiA$_p$-sec24::AFpyrG / pGEM thiA$_p$-sec24::AFriboB*** | |
| sec24 5' SphI F | CGCGGCATGCCGGCTCAAACAAGTCGAAGACCTTATC |
| sec24 5' SpeI R | CGCGACTAGTCTAGGCATTTGCAGCTTATGAACGTTTC |
| sec24 3' SpeI F | CGCGACTAGTATGGCATCTCCACAAGGGGGCTAC |
| sec24 3' NotI R | CGCGGCGGCCGCCACAGACACTTGTGCCTTGGAGCAC |
| AFpyrG SpeI F | CGCGACTAGTGCCTCAAACAATGCTCTTCACCCTC |
| AFriboB SpeI F | CGCGACTAGTCCCGGGCTGCAGGAATTCG |
| thiA$_p$ SpeI R | CGCGACTAGTGTTGACTCAGTTCAATGGTTCGAC |
| ***pGEM thiA$_p$-rabA::AFpyroA*** | |
| rabA 5' AatII F | CGCGGACGTCCGATCGACACAATCGTGCCGTTC |
| rabO 5' SpeI R | CGCGACTAGTCAGGCAGCAGACAAATGGAGAGG |
| rabO ORF SpeI F | CGCGACTAGTATGTCTGAATCAACGCCCGCAAATG |
| rabO 3' NotI R | CGCGGCGGCCGCGTAGCCTTCAGGTCCTGTGTGTTGC |
| AFpyrG SpeI F | CGCGACTAGTGCCTCAAACAATGCTCTTCACCCTC |
| thiA$_p$ SpeI R | CGCGACTAGTGTTGACTCAGTTCAATGGTTCGAC |
| ***pGEM mRFP-rabE::AFpyrG*** | |
| rabE 5' ApaI F | CGCGGGGCCCGAGTGCGGAATATGCCTCCACCTG |
| rabE 5' SpeI R | CGCGACTAGTAGCGAACAGTTAGATACACCGAGGG |
| rabE ORF SpeI F | CGCGACTAGTATGGCTAACGACGAGTATGATGTGAG |
| rabE ORF SpeI R | CGCGACTAGTTTAACAGCATCCACCCTTGTTCTCGG |
| rabE 3' SpeI F | CGCGACTAGTCGTCAACAACGATTTGCGGTTCTG |
| rabE 3' NotI R | CGCGGCGGCCGCCTGTCCAGACCAAAGACCTCCGG |
| mRFP XbaI F | CGCGTCTAGAATGGCCTCCTCCGAGGACGTC |
| mRFP SpeI NS R | CGCGACTAGTGGCGCCGGTGGAGTGGCGG |
| AFpyrG SpeI F | CGCGACTAGTGCCTCAAACAATGCTCTTCACCCTC |
| AFpyrG XbaI R | CGCGTCTAGAACTGTCTGAGAGGAGGCACTGATGCG |
| **pGEM chs5Δ::AFpyrG** | |
| chs5 5' ApaI F | CGCGGGGCCCGAGCCAGAGGCGAAGTTGTGCGGC |
| chs5 5' XbaI R | CGCGTCTAGATCTGACGGAAGCACGAATCGACAG |
| chs5 3' XbaI F | CGCGTCTAGACTGTTGAGCGCTACGTTCGAGTAC |
| chs5 3' NotI R | CGCGGCGGCCGCGGCTCTCCAACCCCTCTTGGGCAAAC |
| AFpyrG XbaI F | CGCGTCTAGAGCCTCAAACAATGCTCTTCACCCTC |
| AFpyrG XbaI R | CGCGTCTAGAACTGTCTGAGAGGAGGCACTGATGCG |
| **pGEM ggAΔ::AFpyrG** | |
| ggA 5' ApaI F | CGCGGGGCCCCTTGGGTACGCCCCTTATCCCTC |
| ggA 5' SpeI R | CGCGACTAGTGACGGCAAAGAGACAGGCTCCTC |
| ggA 3' SpeI F | CGCGACTAGTCGCCACTCACTCGGCCACGATTAG |
| ggA 3' NotI R | CGCGGCGGCCGCCCAGAGCGTTCCAATTCCACCCAG |
| AFpyrG SpeI F | CGCGACTAGTGCCTCAAACAATGCTCTTCACCCTC |
| AFpyrG SpeI R | CGCGACTAGTACTGTCTGAGAGGAGGCACTGATGCG |
| ***pGEM-alcA$_p$-azgA-GFP / pGEM-alcA$_p$-furA-GFP*** | |
| azgA ORF SpeI F | CGCGACTAGTATGGGGTACGCTGAGTGGATTGG |
| furA ORF SpeI F | CGCGACTAGTATGTCAGCTATTAAACGATGGATC |
| GFP NotI R | CGCGGCGGCCGCTTACTTGTACAGCTCGTCCATG |

Gene cassettes were generated by one step ligations or sequential cloning of the relevant fragments in plasmids pBluescript SK II, or pGEM using oligonucleotides carrying additional restriction sites. These plasmids were used as templates to amplify the relevant linear cassettes by PCR. The *thiA$_p$* promoter and a modified version carrying also the FLAG epitope, both fused with the auxotrophic marker *AFpyrG*, were amplified from plasmids pGEM-*thiA*$_p$-*claL* and pGEM-*thiA*$_p$-*FLAG-ap1*$^{\sigma}$, respectively (15). All other gene fusions tagged with fluorescent epitopes carry a 5x Gly-Ala (5xGA) linker, amplified together with GFP or mRFP and *AFpyrG* from plasmids p1439, or p1491 respectively (54). For the construction of *GFP-chsB*, the auxotrophic marker was inserted in a native *Sph*I site at position (-) 1311 from the translation initiation codon. Gene fusions of *alcA*$_p$-*azgA*-GFP and *alcA*$_p$-*furA*-GFP were constructed by cloning the GFP fusions in a modified version of plasmid pGEM-*gpdA*$_p$$^{1000}$-*panB* (49; 53), where the *gpdA*$_p$ promoter fragment was replaced by *alcA*$_p$.

**Movie S1.**
Video showing the exocytosis of *de novo* made UapA-GFP (*$alcA_p$-uapA*, 120 min derepression).

**Movie S2.**
Video showing the co-localization of *de novo* made UapA-GFP (*$alcA_p$-uapA*, 120 min derepression) with sec24-labelled ER-exit sites.
